# Chargaff’s second parity rule lies at the origin of additive genetic interactions in quantitative traits to make natural selection possible

**DOI:** 10.1101/2023.01.19.524724

**Authors:** Bakhyt T. Matkarimov, Murat K. Saparbaev

## Abstract

A major challenge of modern biology is to understand how random genetic variation causes phenotypic variation in quantitative traits. Chargaff’s Second Parity Rule (CSPR) is a statistical property of cellular genomes defined as near exact equalities G≈C and A≈T within single DNA strand. Analysis of mutation spectra inferred from single nucleotide polymorphisms (SNPs) observed in human and mice populations reveals near exact equalities of the reverse complementary mutations, indicating that genetic variations obey CSPR. Furthermore, the nucleotide compositions of coding sequences are statistically interwoven via CSPR since pyrimidine bias at 3^rd^ codon position compensates purine bias at 1^st^ and 2^nd^ positions. Due to this inter-dependence both synonymous and non-synonymous mutations are subject to natural selection. Based on the Fisher’s infinitesimal model, we propose that accumulation of a sufficient number of the reverse complementary mutations results in a continuous trait variation due to small additive effects of statistically inter-linked genetic alterations. Therefore, additive genetic interactions can be inferred as the statistical entanglement of nucleotide compositions of different genetic loci. CSPR challenges the neutral theory of molecular evolution, because all random mutations participate in variation of the traits. The sequence of a gene is interwoven with many other loci in a genome, each of which making infinitely small contribution to a trait variation in contactless manner.

Intra-strand DNA symmetry and genetic variations in cellular organisms. Chargaff’s Second Parity Rule (CSPR), also referred to as intra-strand DNA symmetry, was incidentally determined by measuring nucleotide composition of single-stranded genomic DNAs^1^ and later rediscovered and confirmed by others^2-7^. Noteworthy, intra-strand DNA symmetry can be extended to any pair of reverse complementary short oligonucleotides for a sufficiently long DNA sequence (>100 kb), it means the near exact equalities of reverse complementary pairs of mono- and oligo-nucleotides in one strand of cellular DNA^2,8,9^. CSPR holds for genomes of cellular organisms (including eukaryotic, bacterial and archaeal chromosomes) and double-stranded DNA viruses, whereas, organellar genomes (mitochondria and plastids) smaller than ∼20-30 kbp, single-stranded viral DNA genomes and any type of RNA genome do not obey CSPR^10^. Intriguingly, intra-strand symmetry is maintained in genomic DNA of all cellular organisms despite billions years of divergent evolution of the species from one common ancestor. When regarding at the scale of a chromosome or entire genome, cellular organisms obey CSPR with small deviations less than 1% from mononucleotide parity rule^10^. It remains unknown: what are the mechanisms that force cellular genomes to adapt CSPR and why do genomes of certain non-cellular organisms have broken intra-strand DNA symmetry. Previously it has been proposed that the skewed nucleotide composition of vertebrate mitochondrial genomes is due to asymmetric DNA strand inheritance^11^.

Here, we sought to identify a biological role of intra-strand DNA symmetry in cellular organisms for this we explore a link between CSPR and continuous variation observed in complex traits of higher eukaryotes. Quantitative or complex traits in plants and animals are characterized by measurable and continuous phenotypic variation within a population due to underlying genetic variability and environmental influence. Complex traits such as human height do not follow simple Mendelian inheritance patterns, since their phenotypic variations cannot be explained by the segregation of a single genetic factor. In 1919 Ronald Fisher showed that the variation in continuous traits could be explained by Mendelian inheritance laws if an infinite number of genetic loci control the trait and that each locus contributes additively to the trait^12^. In his work, Fisher introduced the infinitesimal model, also known as polygenic model, based on the idea that variation in a quantitative trait is influenced by an infinitely large number of genes, each of which with an infinitely small (infinitesimal) effect on the phenotype. Importantly, quantitative traits are far more common as compared to simple Mendelian traits; moreover, even the monogenic traits can exhibit phenotypic heterogeneity in organisms with the same genotype because of complex genetic interactions such as incomplete penetrance, variable expressivity and pleiotropy. Since, quantitative traits can be extremely polygenic, they are examined using statistical techniques such as genome-wide association studies (GWAS) rather than classical molecular biology methods^13^. Infinitesimal model provides a basis for statistical modelling and selection theory, which are used in plant and animal breeding programs with great success. Despite this, the model remains a theoretical abstraction, since in reality all living organisms have finite size genomes each containing quite limited number of genes. Prior to the high-throughput DNA sequencing era, it was impossible to verify the infinitesimal model for a quantitative trait in a genetically heterogeneous population. Variants discovered through GWAS accounted for only a small percentage of predicted heritability and this led to the missing heritability problem^14^. With advances in DNA genotyping technologies the problem was in part resolved by demonstrating that most of the heritability could be accounted for by common single-nucleotide polymorphisms (SNPs) missed by GWAS, because their effect sizes fell below genome-wide statistically significant thresholds^15-17^. Furthermore, it has been demonstrated that for most of complex traits the majority of the SNPs-associated heritability is spread uniformly all over the genome^18-20^. Recent studies estimated that more than 10,000 or even more than 100,000 causal variants contribute to a quantitative trait in general, which is in remarkable consistency with infinitesimal model^20,21^.

While many common variants involved in quantitative traits have been identified, the molecular mechanisms through which they contribute to phenotypic variation remain poorly understood^22^. For example, the infinitesimal model states that each genetic locus contributes additively to the trait, while it remains unclear how genetic interactions can act in additive manner and what are the molecular mechanisms of this complementarity. Recently, Boyle and colleagues when analyzing GWAS datasets of the complex traits including human height, schizophrenia, rheumatoid arthritis and Crohn’s disease found that the SNPs associated with traits were evenly distributed all over the genome, rather than clustered within the trait-specific genes^23^. To explain these observations, they proposed the concepts of “core” genes (limited number of disease-specific genes with established biological roles) and “omnigenic” model in which majority of active genes, rather than a specific set of core genes, can affect every complex traits. For example, the total set of expressed genes in an affected tissue or a cell in general outnumber “core” genes by 100:1 or more, consequently the sum of small effects across peripheral genes can far exceed the genetic contribution of variants of core genes because of the highly interconnected cell regulatory networks^23,24^. In support to this, GWAS signals for quantitative traits and disease-associated SNPs are enriched in predicted gene regulatory elements and in open chromatin that is active in cell types relevant to disease, respectively^25-28^.

High polygenic character of quantitative traits and common diseases might also be explained by negative selection towards large effect mutations in the critical genes and loci that play biologically important functions. In support to this idea, statistical modelling of 33 complex traits reveals that heritability is spread more evenly among common variants than among SNPs in functionally important regions due to selective constraints^20^. While researchers were able to identify many genetic factors, such as constitutively active peripheral genes, involved in complex traits, determining the specific molecular mechanisms, through which they contribute to phenotypic variation, remains a major challenge^24^.

## Results and Discussion

### Continuous variation in complex traits enables natural selection

Quantitative traits exhibit continuous normal distribution of phenotypic variations because of their polygenic, multi-locus nature. According to the infinitesimal model proposed by Fisher, continuous phenotypic variation of a quantitative trait, influenced by infinitely large number of genetic loci in a population, have normal distribution (Fig. 1a)^29^. Here we suggest that this bell-shaped distribution also reflects differential selective pressure or fitness of each individual in the population. For example, when considering human height, individuals with heights close to average experience less selective pressure and present in higher numbers in the population since they are better adapted, whereas shortest and tallest individuals endure high selective pressure and thus present in much lower numbers. Similar in multicellular organism, population of somatic cells with mean variations of a trait undergo less selective pressure and present in higher numbers in a given tissue or organ since they are better fit to an internal body environment, whereas cells with extreme phenotypes undergo high selective pressure and present in much lower frequencies because they are less fit. However, as shown in Fig. 1b, variations in the external environment and organism homeostasis can change the direction of selection pressure and as a consequence organisms and somatic cells, respectively, exhibiting positive (+) trait variation would have advantage and this would shift common variants or allele frequency distribution to the right of the abscissa axis.

**Figure 1.**
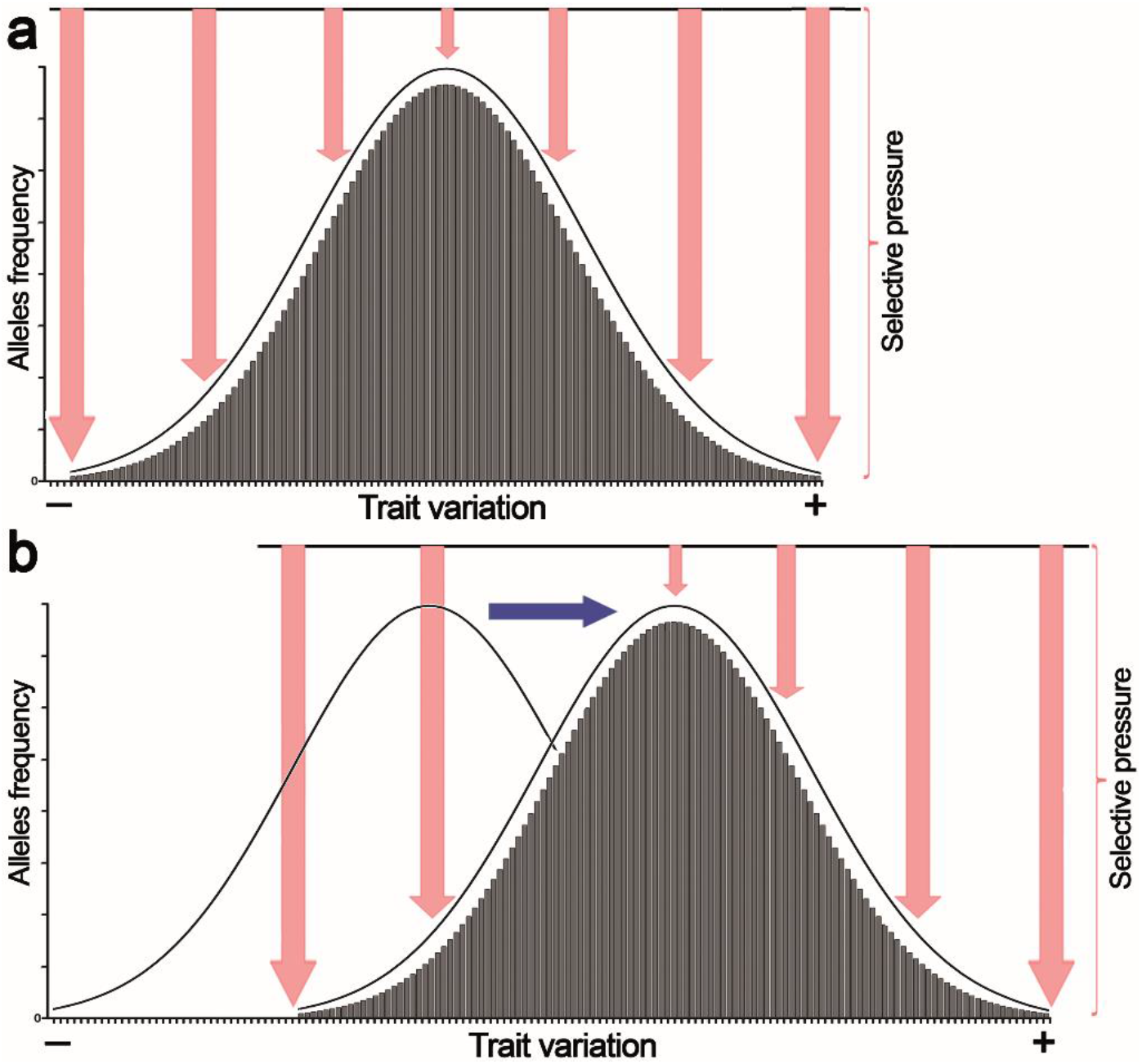
Effect of selective pressure on the normal distribution of a quantitative trait values in a population. (**a**) Stabilizing selection favors the average individuals in a population. (**b**) Directional selection shifts distribution of allele frequencies or genetic variations in a population to adjust trait values to environmental changes. Pink arrows denote selective pressure; blue arrow denotes directional selection.

Cell as a part of highly complex multicellular organism is experiencing a constant selective pressure, because of the internal environment within a body that impose severe restraints on the somatic cells. Multiple factors, among many other such as limited availability of oxygen and nutrients, immune system surveillance, cell-cell contacts, cell-signaling factors that tightly control growth and proliferation of individual cells within a body, all together create selective pressure on a population of somatic cells in order to integrate every part and function of the body as a coherent whole. In fact, cell differentiation, in which changes in the gene transcription programs occurs through epigenetic mechanisms without alterations in the primary DNA sequence, can be regarded as a way for a somatic cell to adapt to continuously changing inner body environment of the complex multicellular organisms during their development and aging. On the other side, random somatic mutations that inevitably accumulate after each cell division, are subject to the strong negative selection pressure that eliminates cells with deleterious and cancer prone mutations. Somatic mutagenesis is a random, stochastic process and majority of mutations observed in higher eukaryotes occur in non-coding regions of the genome and thus may evade the strong selective pressure. It is generally agreed that in cancer only a few (driver) mutations strongly influence its progression, with the remaining (passenger) mutations being neutral or slightly deleterious^30^. Recent Pan-Cancer Analysis of Whole Genomes (PCAWG) variant dataset of the 43,778,859 single nucleotide variants (SNVs) reveal only thousands of mutations that can be identified as drivers, with mean of 5 driver mutations per tumor, whereas, the remaining 99.99% of SNVs have been referred as passenger variants, with no known function and effect on the fitness^31^. Nevertheless, another analysis of PCAWG variant dataset suggest that combined effect of passenger mutations together with undetected weak drivers, can provide additional 12% effect for identifying cancer phenotypes, beyond driver mutations^32^.

Despite huge technological progress in DNA sequencing and analysis of big data, yet fundamental gaps remain in our understanding of how normal cells evolve into tumor cells. It is assumed that the absolute majority of somatic and passenger mutations in normal and cancer cells, respectively, are either neutral or slightly deleterious. However, this assumption is challenged by several important observations. The existence of genetic interaction networks between multiple genes on a genome-wide scale that can determine phenotypic variability of the observable traits including the expression of monogenic traits such as rare, highly penetrant, deleterious gene mutations. It has been shown that common genetic variants can critically contribute to incomplete penetrance of severe Mendelian diseases, and that not all individuals with a given genotype (mutant allele) display a corresponding phenotype^33^. This phenomenon was observed also on other model organisms (reviewed in^34^). Numerous studies have convincingly demonstrated that Calorie Restriction (CR), as compared to control Ad Libitum (AL) regimen, dramatically delays spontaneous tumor development in laboratory animals^35^. This dietary regimen changes the body’s homeostasis in a way that might reduce the fitness of cancer-prone cells as compared to normal cells and prevents their propagations in normal tissues. Here, we propose that accumulation of sufficient number of random mutations in somatic cells inevitably results in the phenotypic variations owing to small additive genetic interactions. Trait variations in somatic cells such as glucose uptake and proliferation rate are subject to natural selection. Therefore, CR regimen would favor the growth of cells with lower proliferation rate and glucose consumption, whereas, pre-cancerous cells would be under increased negative selection pressure and this in turn would prevent their propagation. Thus, we suggest that common variants and passenger mutations can influence the phenotypic expression of a simple Mendelian trait such as a cancer-driver mutation by uncovering its complex polygenic character. Below, we will analyze the role of intra-strand DNA symmetry in genetic variations and phenotypic diversity.

### Intra-strand DNA symmetry and genetic variations in cellular organisms

CSPR can be interpreted as the law of large numbers which states that, as the number of identically distributed, randomly generated variables increases, their sample mean approaches their distribution mean. Therefore, intra-strand DNA symmetry can be considered as a general statistical property of cellular genomes composed of independently segregating replicons which is maintained over the evolutionary history of life. This means that a sufficiently large number of genetic variations in a population have a statistical tendency in that every single random mutation must be “*compensated*” by another inverse complementary mutation elsewhere in the genome. This mechanism of compensation manifests itself through the effect of paired reverse-complementary mutations (e. g. C→T pairs with G→A) occurring in one DNA strand that enables to maintain the intra-strand DNA symmetry within entire segregating replicon(s). As shown in Table 1, analysis of the patterns of spontaneous mutations in trinucleotide sequence context inferred from single nucleotide polymorphisms (SNPs) observed in human and mice populations revealed that in single DNA strand of human and mice genomes, the number (V) of X↔Y variations in triplet NXN (referred to here as V_NXN↔NYN_) is nearly equal to the number of X’↔Y’ variations occurring in reverse complementary triplet N’X’N’ on the same DNA strand (referred to here as V_N’X’N’↔N’Y’N’_, where, V is for total number of a variation, N is any nucleotide, X and Y are different versions of a nucleotide, N’, X’ and Y’ are corresponding reverse complementary counterparts). For example, the number of SNP AGT↔AAT, referred to here as V_AGT↔AAT_, observed in human population, is 2.375.298 (100%), which is nearly equal to the number of SNP of its reverse complementary version V_ACT↔ATT_, observed 2.367.625 times (99.7%). Similar statistical parities are observed in mice populations there V_AGT↔AAT_ ≈ V_ACT↔ATT_ (384.516 (100%) and 384.248 (99.99%) times, respectively) with less than 1% difference (Table 1 and S1).

**Table 1.** Mutation spectra inferred from single nucleotide polymorphisms in human and mice populations.

| N° | N X N | N Y N | X | Y | Total number of SNPs | Fraction | N'X'N' | N'Y'N' | X' | Y' | Total number of SNPs | Fraction | Difference (%) |
| --- | --- | --- | --- | --- | --- | --- | --- | --- | --- | --- | --- | --- | --- |
| **Human SNPs** |  |  |  |  |  |  |  |  |  |  |  |  |  |
| 1 | AGC | ATC | G | T | 312720 | 0.37 | GAT | GCT | A | C | 310244 | 0.36 | 0.79 |
| 2 | CAA | CTA | A | T | 227826 | 0.27 | TAG | TTG | A | T | 229630 | 0.27 | 0.79 |
| 3 | ACT | ATT | C | T | 2367625 | 2.77 | AAT | AGT | A | G | 2375298 | 2.78 | 0.32 |
| 4 | AAA | AGA | A | G | 1729945 | 2.02 | TCT | TTT | C | T | 1734336 | 2.03 | 0.25 |
| 5 | TAT | TTT | A | T | 544308 | 0.64 | AAA | ATA | A | T | 543030 | 0.64 | 0.23 |
| 6 | GAG | GTG | A | T | 287742 | 0.34 | CAC | CTC | A | T | 288408 | 0.34 | 0.23 |
| 7 | GCG | GTG | C | T | 2008008 | 2.35 | CAC | CGC | A | G | 2008195 | 2.35 | 0.01 |
| 8 | TAA | TGA | A | G | 1448254 | 1.69 | TCA | TTA | C | T | 1448196 | 1.69 | 0.00 |
| **Mouse SNPs** |  |  |  |  |  |  |  |  |  |  |  |  |  |
| 1 | CGA | CTA | G | T | 38698 | 0.26 | TAG | TCG | A | C | 38235 | 0.25 | 1.20 |
| 2 | AAG | ATG | A | T | 88927 | 0.59 | CAT | CTT | A | T | 89884 | 0.59 | 1.06 |
| 3 | GAG | GCG | A | C | 50929 | 0.34 | CGC | CTC | G | T | 50724 | 0.33 | 0.40 |
| 4 | CAG | CCG | A | C | 56636 | 0.37 | CGG | CTG | G | T | 56410 | 0.37 | 0.40 |
| 5 | GAT | GTT | A | T | 63675 | 0.42 | AAC | ATC | A | T | 63432 | 0.42 | 0.38 |
| 6 | ACT | ATT | C | T | 384248 | 2.53 | AAT | AGT | A | G | 384516 | 2.54 | 0.07 |
| 7 | TAT | TTT | A | T | 103335 | 0.68 | AAA | ATA | A | T | 103323 | 0.68 | 0.01 |
| 8 | AAG | AGG | A | G | 299963 | 1.98 | CCT | CTT | C | T | 299979 | 1.98 | 0.01 |

Furthermore, analysis of small insertion/deletion (indel) polymorphism in human population revealed near exact equalities in the number of indels between 18 out of 32 reverse complementary pair of trinucleotides (Table S2). These trinucleotides have sequence context in which a base corresponding to indel is different from the neighboring base (e.g. V_TΔC↔TGC_ ≈ V_GΔA↔GCA_). It should be noted that for 14 out of 32 pairs, the correct pairwise alignment was not possible because of sequence context particularity. These trinucleotides have sequence context in which a base corresponding to indel is the same as the neighboring base (e.g. V_TΔG↔TTG_ ≠ V_CΔA↔CAA_). These observations indicate that during evolution the accumulation of random mutations in a population of cellular organisms, represented here in the form of SNPs and indels, is necessary and sufficient condition to drive and maintain chromosomal DNA in the state of intra-strand symmetry equilibrium. Thus, the stochastic processes such as random mutations can lead genome evolution towards the symmetry of nucleotide compositions and this should be considered as one of the major force of evolution, which provides great selective advantage to cellular organisms.

### Statistical inter-dependence of the nucleotide compositions is revealed by a bias at 3^rd^ position of codon

The analysis of mutation spectra inferred from SNPs suggest that CSPR is an essential property of cellular organisms, which restrains the patterns of random genetic mutations to make them assembled in the pairwise, statistically interwoven manner. Local deviations from intra-strand DNA symmetry in the majority of coding DNA sequences (CDSs), referred as to Szybalski’s rule^36^ (when purines exceed pyrimidines), suggest that at the gene level, directional mutation pressures and selective constraints counteracts the homogenizing force of CSPR. To examine this closely, we measured the deviations from intra-strand DNA symmetry of the nucleotide composition at each of three positions of a codon in bacteria and human CDSs and presented data in the graph in which each codon position is plotted according to its purine/pyrimidine (Pur/Pyr or G/C and A/T) ratio. As shown in Fig. 2a, when measuring Pur/Pyr ratio for all three positions of a codon together, the nucleotide compositions of each triplet (denoted as black squares) in bacterial proteomes exhibit relatively weak deviations from CSPR with G/C and A/T ratio nearly equal to one (≈ 1), as compared to that of single codon positions (denoted as colored squares). Remarkably in absolute majority of bacteria, 1^st^ position of a codon (red square) has highly enriched purine content (Pur/Pyr ratio ≥ 1.5), while positions 2 and 3 (blue and green squares, respectively) contain increased C and T content (G/C and A/T ratio ≤ 1), respectively, indicating that the nucleotide composition at 2^nd^ and 3^rd^ positions of a codon compensate very strong bias in the purine content at 1^st^ position of a codon. As a result, when all three positions of a codon are taken into account, we observe more symmetry in the nucleotide compositions of triplets, which in turn dramatically reduces the deviations of CDSs from CSPR at a gene level in bacteria. In human proteome we can also observe the phenomenon of compensation of purine bias in the nucleotide composition at 1^st^ position of a codon by that of 2^nd^ and 3^rd^ positions, which results in Pur/Pyr ratio ≈ 1 for the nucleotide composition of each triplet (Fig. 2b). Two last codon positions have enriched pyrimidine content (>50%) which is opposite to that of 1^st^ position of a codon, thus again the Pur/Pyr content at 2^nd^ and 3^rd^ positions compensate the strong bias from intra-strand DNA symmetry of the nucleotide composition at 1^st^ position of a codon in human CDSs.

**Figure 2.**
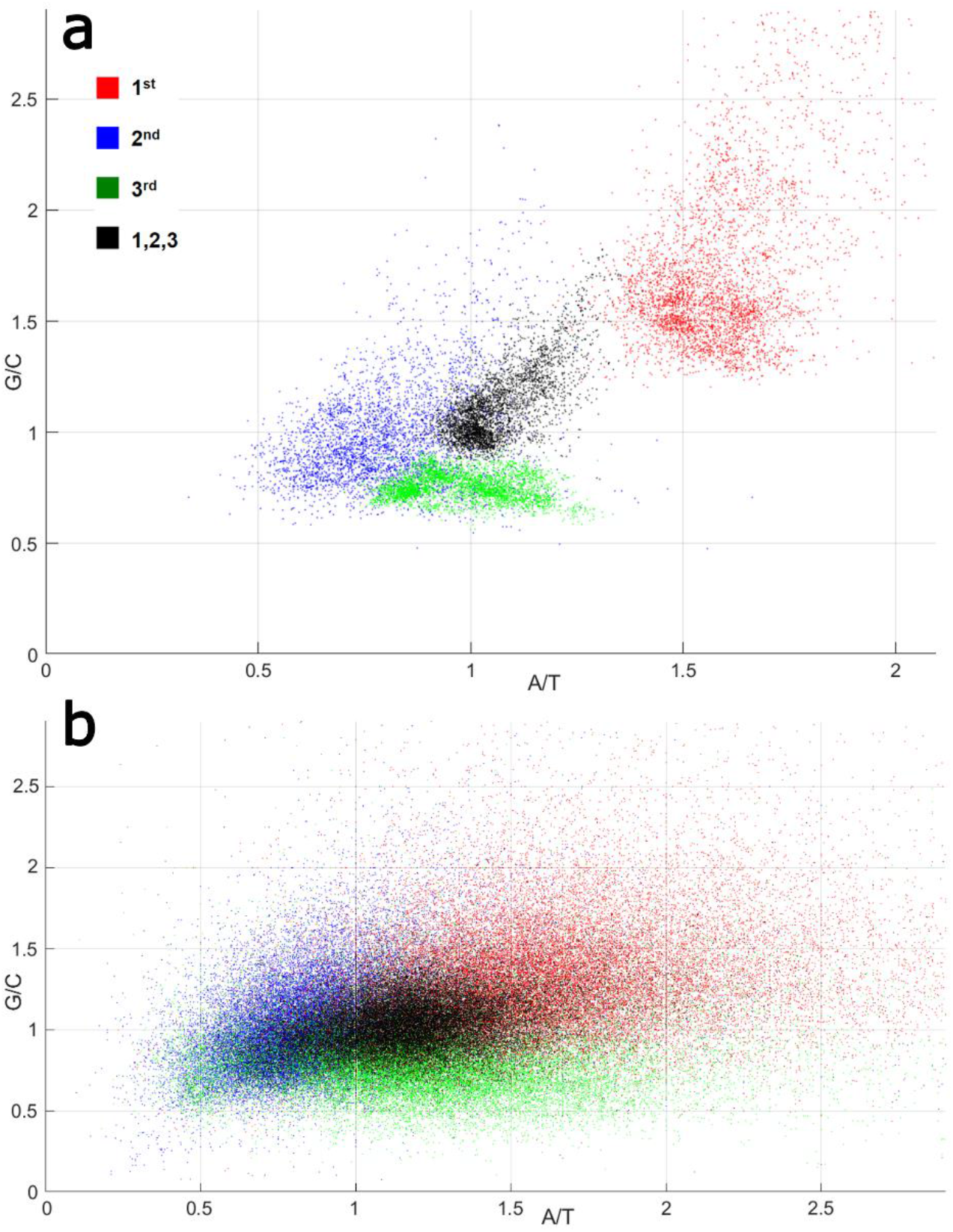
Nucleotide compositions at positions of a codon in bacterial and mammalian proteomes. G/C and A/T ratio for each position of a codon denoted as a red square for 1^st^ position; blue for 2^nd^ position; green for 3^rd^ position; and black for all three positions together. (**a**) Bacterial ORFs. Each point corresponds to a proteome of one bacterial strain. (**b**) Human ORFs. Each points corresponds to a single ORF.

Noteworthy, due to redundancy of the genetic code, mutations at the 3^rd^ position of a codon often results in a synonymous change of amino acid sequence of the protein, which explains lower evolutionary conservation of this position as compared to first two positions of a codon^37^. Mutations at 1^st^ and 2^nd^ positions of a codon in majority of cases lead to non-synonymous changes in the protein sequence, which are subject to strong selection pressure, therefore, the first two positions of a codon are much more functionally constrained, as compared to 3^rd^ one. Nevertheless, our data indicate that variations in the nucleotide composition at 3^rd^ position of a codon exhibit strong statistical entanglement or inter-dependency with that of 1^st^ and 2^nd^ positions, owing to the law of large numbers that maintains CSPR at genome level. *Based on these observations, we suggest that the functional constraints and selective pressure towards first two codon positions, also act on 3*^*rd*^ *codon position, owing to the statistical entanglement of their nucleotide compositions*. As a result, an increase in purine content at 3^rd^ position of codons due to the synonymous transversion mutations would be under strong selection pressure. Indeed, the 3^rd^ codon position has a much higher transition/transversion mutation bias than the 1^st^ and 2^nd^ codon position suggesting that transversions are disproportionally selected against at the last position of a codon^38^. Thus, under intra-strand DNA symmetry equilibrium of a gene sequence, both synonymous and non-synonymous mutations in a CDS would be under selective pressure because of their inter-dependence.

Taken together these observations suggest that the synonymous mutations at 3^rd^ position of a codon, when they accumulate in sufficient number, are neither neutral, nor nearly neutral, but subject to natural selection to the similar extent as the mutations in 1^st^ and 2^nd^ codon positions. Hence, by virtue of CSPR, random mutations occurring in cellular genomes at various genetic loci over generations become statistically interwoven and this makes them all subject to natural selection. Moreover, when considering genetic loci such as junk DNA in higher eukaryotes^39^, which are seemingly useless and have no known function, their nucleotide compositions are statistically interwoven with that of functional regulatory and coding DNA sequences, as a consequence, natural selection can act on the genetic variations in both functional and junk DNA. Owing to CSPR, accumulation of mutations in junk DNA will exert increasing selective pressure on the mutation pattern in functional (regulatory and coding) DNA sequences, forcing natural selection act on seemingly functionless DNA regions.

### Intra-strand DNA symmetry and the origin of additive genetic interactions

CSPR acts at full strength when a sufficiently big number of random mutations are taken into consideration, for example in a large set, the mutations can be assembled as pairs of reverse complementary matches (see Table 1). Whereas small number of random mutations may not always exhibit the expected intra-strand DNA symmetry, nevertheless with each additional mutation the probability that all observed mutations would be in accordance with CSPR increases. Consequently, when the number of mutations occurring in various genetic loci increases in time dependent manner, they would approach intra-strand DNA symmetry and exhibit strong statistical entanglement. For that reason, increasing number of mutations in non-coding DNA and peripheral gene networks should be compensated in part by the reverse complementary mutations in coding DNA and core gene network, respectively, and vice versa, so that the nucleotide compositions of an entire chromosome comply with CSPR. Furthermore, the homogenizing force of CSPR acts also at local gene level, over the course of multiple generations the nucleotide composition of a CDS should be approaching closely the intra-strand DNA symmetry, despite structural and functional constraints on the evolution of a protein (Fig. 2, black squares).

Above analysis of SNPs and biases at different codon positions further demonstrate that intra-strand DNA symmetry is a general property of cellular organisms that underlies statistical inter-dependence of the nucleotide compositions of different genetic loci and patterns of spontaneous mutations. *Hence, we suggest that this statistical entanglement provides a long-searched mechanism of the additive genetic interactions, which constitute the basis of continuous variation in expression of the traits*. According to Fischer’s infinitesimal model, continuous phenotypic variation in a quantitative trait is achieved through additive genetic interactions of an infinitely large number of genetic loci^12^. In general, the effect of a single mutation is infinitesimally small and cannot be measured experimentally. However, the cumulative effect of many such mutations become visible and can be measured, since the mutations become statistically entangled in the course of evolution, and their accumulation give rise to additive genetic effect. *Here, we propose that the additive genetic effect is achieved through intra-strand complementarity of the nucleotide compositions of diverse genetic loci participating in a trait variation*. The nucleotide compositions of coding DNA sequences and regulatory elements (such as promoters and origins of replication) obey CSPR. In higher eukaryotes these highly conserved DNA sequences are statistically interwoven with other sites in a genome, which in their majority are rather non-coding and non-conserved. Consequently, when a sufficient number of the genetic variations in non-coding DNA sequences are accumulated, this would result in continuous variation of a trait via a modulation of the expression of highly conserved DNA sequences.

Because of CSPR, which can be derived from the law of large numbers, random genetic mutations in a cell or an organism have a statistical tendency to accumulate in pairwise manner in which pair of reverse complementary SNPs present in near equal numbers V_X↔Y_ ≈ V_X’↔Y’_ or V_NXN↔NYN_ ≈ V_N’X’N’↔N’Y’N’_ (e.g. the numbers of G↔A or TGC↔TAC SNPs in a population are near equal to that of C↔T or GCA↔GTA SNPs, respectively (V_G↔A_ ≈ V_C↔T_ or V_TGC↔TAC_ ≈ V_GCA↔GTA_)). This means that accumulation of C→T mutations would be accompanied with nearly exact number of G→A mutations in the same DNA strand of a chromosome distributed randomly anywhere in a genome. As shown in Fig. 3, we propose that accumulation of the reverse-complementary mutations results in additive genetic effect, which under directional selection would shift the phenotype towards one end of a trait spectrum.

**Figure 3.**
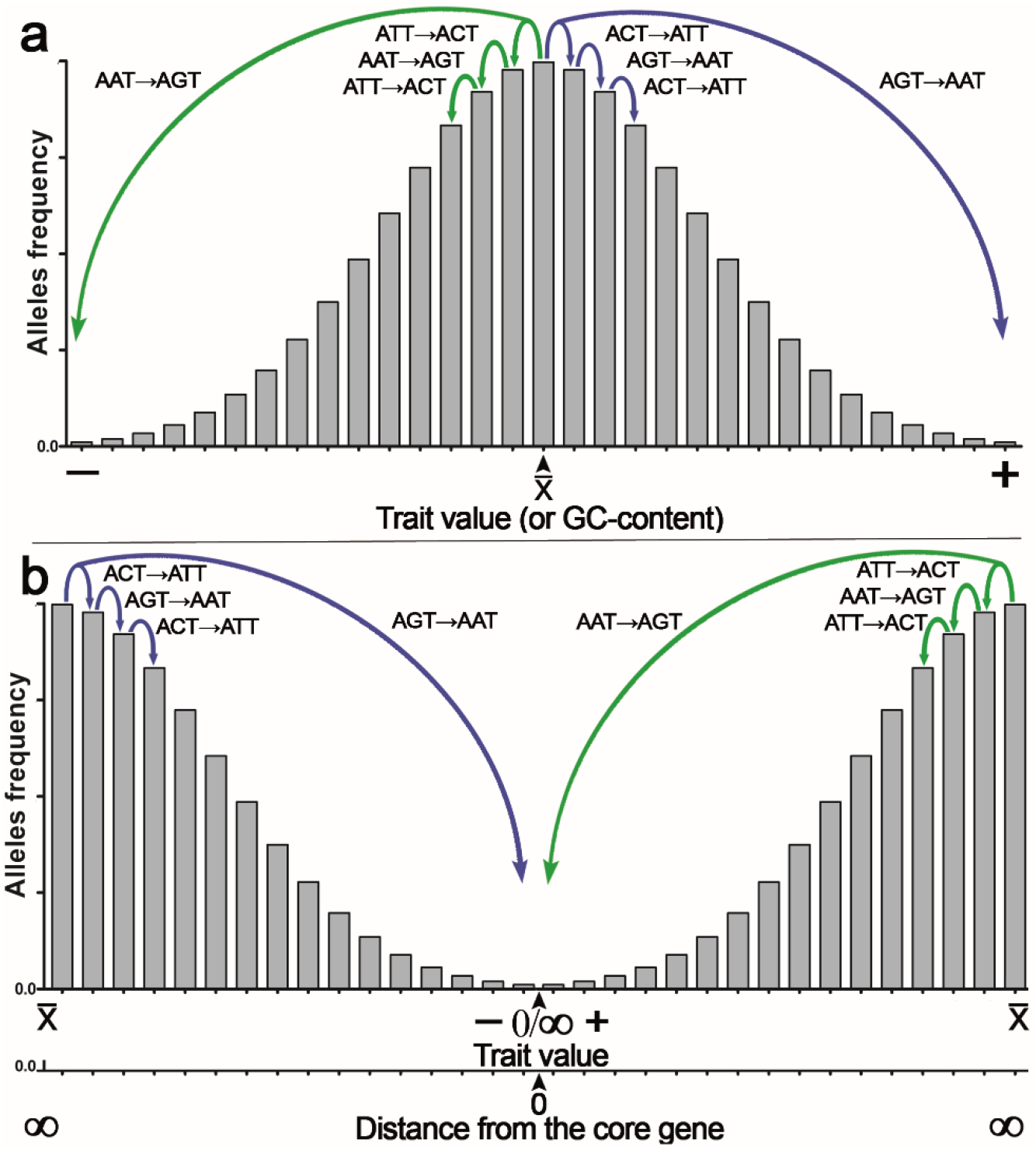
Graphical representation of the additive effect of mutations on variations in the expression of a quantitative trait. (**a**) The normal distribution of quantitative a trait with mean value of a trait 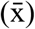 at the centre of X-axis. (**b**) The normal distribution of quantitative a trait with mean value of a trait 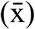 at the extremes of X-axis. Note that the distance from core gene is plotted in additional X-axis at the bottom. Accumulation of the statistically inter-dependent mutations in the CSPR-mode creates continuous trait variation. The majority of pairwise mutations makes an infinitely small additive contribution to the trait variations (e.g. glucose uptake by a somatic cell). Accumulation of “forward” C→T/G→A mutations lead to an increase, whereas “reverse” T→C/A→G mutations lead to a decrease in expression of the trait. Under directional selection the difference between numbers of “forward” and “reverse” mutations increases.

For example, a pairwise accumulation of either “forward” T→C/A→G or “reverse” C→T/G→A mutations in a population of organisms or somatic cells would have increasing additive effect on the expression of a trait. Importantly, accumulation of T→C/A→G mutations would shift a trait variation to the left or “minus” side of the normal distribution (let’s say lower glucose consumption rate of a somatic cell), whereas reverse C→T/G→A mutations would shift trait variation in opposite direction to the right or “plus” side (e. g. higher glucose consumption rate). According to this model some rare mutations if occur in a core gene may have large effect on the expression of a trait, subsequently they would be under strong negative selection^20^. Whereas, the majority of reverse-complementary mutations would have rather small effect, however their effects would be additive and their accumulation under directional selection would lead to continuous variation in the expression of a trait from small to large effect depending on the number of reverse-complementary mutations or SNPs. It is tempting to speculate that the effect size of an SNP would be inversely proportional to the distance between SNP and “core” gene (Fig. 3b).

Thus, accumulation of either T→C/A→G or C→T/G→A mutations in pairwise manner over many generations will give rise to the continuous variations in the expression of a trait in a population and at the same time will maintain intra-strand DNA symmetry. Under directional selection which would favor proliferation of cells or organisms with either decreased (-) or increased (+) expression of the trait, accumulation of the reverse-complementary mutations with additive effect may change the GC content, which is one of the integral feature of genome. For example, GC content would decrease, if due to differential selection the “forward” C→T/G→A and C→A/G→T mutations accumulate at higher rate as compared to their “reverse” counterparts T→C/A→G and A→C/T→G mutations, respectively. Whereas, accumulation of the transversions C→G/G→C and T→A/A→T could modify GC and AT skews, respectively, if they occur on a chromosome in the symmetric positions with respect to the minimum or maximum of a skew.

Analysis of the patterns of common variants affecting human height, the classic example of a quantitative trait, shows that majority of 697 SNPs identified by GWAS are T*↔C (231) and A*↔G (245) (* denotes the effect allele)^27^. We observe that V_T*↔C_ ≈ V_A*↔G_ as expected, and surprisingly no “reverse” C*↔T and G*↔A variants are perceived among other variants. Another example of the additive effect of reverse-complementary mutations is the mutational signatures in cancer, in recent version of COSMIC (the Catalogue Of Somatic Mutations In Cancer) database v3.3 - June 2022 for the single base substitutions (SBS) about 60 different types of mutational signatures are listed^40^. Remarkably, the majority of SBS signatures in cancer 48 out of 60 can produce a change in GC content, for example in SBS1, 2, 6, 7ab, 11, 15, 19, 23, 31 and 32 the number of C→T mutations are much higher than that of reverse T→C mutations, thus accumulation of the former results in very small decrease in GC content of the cancer genomes. It should be noted that the algorithm which generates SBS signatures take in account only 6 possible base substitutions instead of 12, this operation is made by adding together pyrimidine and purine substitutions e.g. C→T and G→A mutations are presented as C→T only, whereas C→A and G→T mutations shown as C→A only, this algorithm cannot tell us whether V_C→T_ ≈ V_G→A_. Finally, accumulation of the reverse complementary T*↔C/A*↔G SNPs in human population and C→T/G→A mutations in certain common cancer types, associated with a minor change in the GC content, support our hypothesis that these genetic variations are additive and thus can shift the trait variation to either end of distribution (Fig. 3).

### Conclusions

The golden standard of genetic studies is to examine the gene function or effect of mutation by employing isogenic strains in order to compare a wild-type strain to a strain that is genetically identical with the exception of the gene or mutation of interest. It is postulated that the use of isogenic strains rules out the effect of genetic background or differences between the genomes of organisms under investigation. Indeed, even subtle differences in distant genetic loci might influence expression of a gene or mutation studied owing to *the genetic interactions*. Thus, this trivial observation indicates that various forms of genetic interactions unify different parts of a genome into a whole. In the infinitesimal model an infinitely large number of genetic loci influence expression of complex trait via additive genetic interactions each of which making an infinitely small, immeasurable contribution to the phenotype. The nature of additive genetic interactions cannot be understood in a pure mechanistic manner, but rather by means of statistical approach. Mechanistic models describe specific biologically relevant functions and deal with a finite number of genes and other genetic elements, which interact in explicit manner to perform their functions, and encoded by a finite number of genetic loci. Whereas, complex traits, such as the size of cells, human height and cancer, deal with a large number of biological functions and associated with them proteins and other macromolecules and, consequently, encoded and regulated by a large number of genetic loci. In that case, interactions between genetic loci occur rather in implicit manner since they are based on the statistical relationship between their nucleotide compositions and genetic variations. Nucleotide composition of 3^rd^ position of a codon compensating purine bias in two first positions provide an example of implicit interaction which we call a statistical entanglement. Here, we suggest that the statistical entanglement is a contactless “mechanism” for additive genetic interactions in the Fisher’s infinitesimal model. Consequently, the nucleotide composition of a gene directly affecting a trait via physical action of the protein that it encodes is statistically interwoven with many different loci in a genome, each of which making an infinitely small additive contribution to the phenotypic variations. Thus, the statistical entanglement of nucleotides in DNA challenges the neutral theory of molecular evolution. For the cellular genomes obeying CSPR there are neither neutral, nor nearly neutral mutations, because of the additive genetic interactions to which all random mutations participate in contact-free manner. In summary, intra-strand DNA symmetry equilibrium of a cellular genome, which takes origin from stochastic processes, generates regulatory, highly interconnected genetic network in living organisms and makes Darwinian evolution possible.

## Methods

### Analysis of mutation spectra derived from SNP variation in natural populations

NCBI dbSNP is a public collection of nucleotide variations for different species from a wide variety of sources^41^. DbSNP do not track individual samples and represent all identified genetic variation aligned to reference genome. SNP/indels statistics were computed on NCBI dbSNP dataset build 142 with reference genomes: human GRCh38 build 38.1, mouse GRCm38.p2 build 38.3. Every SNP counted once from top strand, only validated SNP/indels are counted, SNPs from sex chromosomes are excluded. Every SNP/indel was enriched by left/right flanking nucleotide sequences. There are 192 directed SNP of type NXN↔NYN, where X/Y are ref/alt nucleotides. These 192 SNP’s are grouped by 88 symmetric pairs based on ref/alt interchange, e.g. the count of group of AGC↔ATC was compared with the count of reverse complementary group of GAT↔GCT. In addition there are 8 self-complementary SNPs, which are presented as a single group (e.g. AGT↔ACT, CAG↔CTG …).

### Analysis of the nucleotide composition at different codon positions in bacteria and human CDSs

Protein coding sequences (CDS) were selected from the curated databases, CCDS for human/mouse^42^ and NCBI RefSeq for bacterial specie^43^. For bacterial genomes we exclude CDSs for hypothetical proteins. For every selected CDS we counted statistics of each single nucleotide for 1^st^, 2^nd^ and 3^d^ codon positions, as well as CDS total statistics, and then compute corresponding A/T and G/C ratios, as follows we have four (A/T, G/C) ratios that includes the ratios for three different positions plus ratio for all three positions together. For human CDS we have created two dimensional plot of (A/T, G/C) for every CDS separately, in total 33420 CDS, with different colors for codon positions, and black color for CDS total. For bacteria we computed accumulative statistics, e.g. single nucleotide counts are sum of all CDSs from single bacterial species, this produces four (A/T, G/C) ratios for each species separately, in total 4038 species that contains 60417 different names of proteins.

## ABBREVIATIONS

CSPR: Chargaff’s Second Parity Rule
SNP: single nucleotide polymorphism
GWAS: genome-wide association studies
V: total number of SNPs or mutations.

## Acknowledgments

This work was supported by the Science Committee of the Ministry of Education and Science of the Republic of Kazakhstan (grant AP09260233) and Nazarbayev University CRP (grant 091019CRP2111) to B.T.M.; and by the French National Research Agency (grant ANR-22 CE1212-0034-01) and Electricité de France (grants RB 2020-02 and RB 2021-05) to M.K.S.

## Author contributions

B.T.M. and M.K.S. designed the study. B.T.M. conceived of the methodology and data collection. M.K.S. wrote the manuscript. All authors discussed and contributed to the analysis of data and to the final version of the paper.

## Competing interests

The authors declare no competing interests.

## Data availability

Data sharing not applicable—no new data are generated.

## Notes

### Competing Interest Statement

The authors have declared no competing interest.

