## Supplemental Tables S1-2 for "Chargaff’s second parity rule lies at the origin of additive genetic interactions in quantitative traits to make natural selection possible"

**Supplementary Table S1**. Mutation spectra inferred from single nucleotide polymorphisms in human and mice populations.

| N° | N X N | N Y N | X | Y | Total number of SNPs | Fraction | N'X'N' | N'Y'N' | X' | Y' | Total number of SNPs | Fraction | Difference (%) |
|---|---|---|---|---|---|---|---|---|---|---|---|---|---|
| | **Human SNPs** | | | | | | | | | | | | |
| 1 | AGC | ATC | G | T | 312720 | 0.37 | GAT | GCT | A | C | 310244 | 0.36 | 0.79 |
| 2 | CAA | CTA | A | T | 227826 | 0.27 | TAG | TTG | A | T | 229630 | 0.27 | 0.79 |
| 3 | TCT | TGT | C | G | 902323 | 1.06 | ACA | AGA | C | G | 895602 | 1.05 | 0.74 |
| 4 | CGT | CTT | G | T | 296420 | 0.35 | AAG | ACG | A | C | 294421 | 0.34 | 0.67 |
| 5 | TAT | TGT | A | G | 3028959 | 3.54 | ACA | ATA | C | T | 3010412 | 3.52 | 0.61 |
| 6 | AGT | ATT | G | T | 472582 | 0.55 | AAT | ACT | A | C | 469735 | 0.55 | 0.60 |
| 7 | CCT | CGT | C | G | 379974 | 0.44 | ACG | AGG | C | G | 377686 | 0.44 | 0.60 |
| 8 | CCC | CGC | C | G | 306967 | 0.36 | GCG | GGG | C | G | 305208 | 0.36 | 0.57 |
| 9 | GAT | GTT | A | T | 427135 | 0.50 | AAC | ATC | A | T | 424703 | 0.50 | 0.57 |
| 10 | AAA | ACA | A | C | 887037 | 1.04 | TGT | TTT | G | T | 891542 | 1.04 | 0.51 |
| 11 | GAC | GCC | A | C | 377851 | 0.44 | GGC | GTC | G | T | 379592 | 0.44 | 0.46 |
| 12 | TGA | TTA | G | T | 494282 | 0.58 | TAA | TCA | A | C | 496509 | 0.58 | 0.45 |
| 13 | AAG | ATG | A | T | 441016 | 0.52 | CAT | CTT | A | T | 442585 | 0.52 | 0.35 |
| 14 | ACT | ATT | C | T | 2367625 | 2.77 | AAT | AGT | A | G | 2375298 | 2.78 | 0.32 |
| 15 | GAC | GGC | A | G | 1039990 | 1.22 | GCC | GTC | C | T | 1036758 | 1.21 | 0.31 |
| 16 | ACG | ATG | C | T | 3502135 | 4.10 | CAT | CGT | A | G | 3511739 | 4.11 | 0.27 |
| 17 | TCC | TTC | C | T | 1292114 | 1.51 | GAA | GGA | A | G | 1288797 | 1.51 | 0.26 |
| 18 | AAA | AGA | A | G | 1729945 | 2.02 | TCT | TTT | C | T | 1734336 | 2.03 | 0.25 |
| 19 | TAT | TTT | A | T | 544308 | 0.64 | AAA | ATA | A | T | 543030 | 0.64 | 0.23 |
| 20 | GAG | GTG | A | T | 287742 | 0.34 | CAC | CTC | A | T | 288408 | 0.34 | 0.23 |
| 21 | GAA | GCA | A | C | 512618 | 0.60 | TGC | TTC | G | T | 513797 | 0.60 | 0.23 |
| 22 | ACC | ATC | C | T | 1331428 | 1.56 | GAT | GGT | A | G | 1334457 | 1.56 | 0.23 |
| 23 | GCA | GTA | C | T | 1234387 | 1.44 | TAC | TGC | A | G | 1237171 | 1.45 | 0.23 |
| 24 | GAG | GCG | A | C | 240273 | 0.28 | CGC | CTC | G | T | 240790 | 0.28 | 0.21 |
| 25 | TCC | TGC | C | G | 444969 | 0.52 | GCA | GGA | C | G | 444029 | 0.52 | 0.21 |
| 26 | GGG | GTG | G | T | 467854 | 0.55 | CAC | CCC | A | C | 468761 | 0.55 | 0.19 |
| 27 | GGA | GTA | G | T | 367737 | 0.43 | TAC | TCC | A | C | 368387 | 0.43 | 0.18 |
| 28 | TGG | TTG | G | T | 543832 | 0.64 | CAA | CCA | A | C | 542895 | 0.64 | 0.17 |
| 29 | CAA | CGA | A | G | 1798724 | 2.10 | TCG | TTG | C | T | 1801266 | 2.11 | 0.14 |
| 30 | CGA | CTA | G | T | 160741 | 0.19 | TAG | TCG | A | C | 160957 | 0.19 | 0.13 |
| 31 | TCG | TGG | C | G | 280491 | 0.33 | CCA | CGA | C | G | 280147 | 0.33 | 0.12 |
| 32 | GCT | GTT | C | T | 1378880 | 1.61 | AAC | AGC | A | G | 1380364 | 1.62 | 0.11 |
| 33 | CAG | CCG | A | C | 292541 | 0.34 | CGG | CTG | G | T | 292773 | 0.34 | 0.08 |
| 34 | TAC | TTC | A | T | 230424 | 0.27 | GAA | GTA | A | T | 230244 | 0.27 | 0.08 |

| | | | | | | | | | | | | | |
|---|---|---|---|---|---|---|---|---|---|---|---|---|---|
| 35 | AGA | ATA | G | T | 604514 | 0.71 | TAT | TCT | A | C | 604874 | 0.71 | 0.06 |
| 36 | CCG | CTG | C | T | 2588225 | 3.03 | CAG | CGG | A | G | 2589637 | 3.03 | 0.05 |
| 37 | GGT | GTT | G | T | 611030 | 0.71 | AAC | ACC | A | C | 611362 | 0.72 | 0.05 |
| 38 | AAG | AGG | A | G | 1664646 | 1.95 | CCT | CTT | C | T | 1663799 | 1.95 | 0.05 |
| 39 | CAT | CCT | A | C | 481839 | 0.56 | AGG | ATG | G | T | 482018 | 0.56 | 0.04 |
| 40 | CCA | CTA | C | T | 1383378 | 1.62 | TAG | TGG | A | G | 1383081 | 1.62 | 0.02 |
| 41 | GCT | GGT | C | G | 418689 | 0.49 | ACC | AGC | C | G | 418602 | 0.49 | 0.02 |
| 42 | GAG | GGG | A | G | 1195336 | 1.40 | CCC | CTC | C | T | 1195523 | 1.40 | 0.02 |
| 43 | GCG | GTG | C | T | 2008008 | 2.35 | CAC | CGC | A | G | 2008195 | 2.35 | 0.01 |
| 44 | TAA | TGA | A | G | 1448254 | 1.69 | TCA | TTA | C | T | 1448196 | 1.69 | 0.00 |
| 45 | TAA | TTA | A | T | 532422 | 0.62 | | | | | | | |
| 46 | CAG | CTG | A | T | 343352 | 0.40 | | | | | | | |
| 47 | GCC | GGC | C | G | 364797 | 0.43 | | | | | | | |
| 48 | ACT | AGT | C | G | 690362 | 0.81 | | | | | | | |
| 49 | CCG | CGG | C | G | 135675 | 0.16 | | | | | | | |
| 50 | AAT | ATT | A | T | 533699 | 0.62 | | | | | | | |
| 51 | TCA | TGA | C | G | 596062 | 0.70 | | | | | | | |
| 52 | GAC | GTC | A | T | 256117 | 0.30 | | | | | | | |
| N° | N X N | N Y N | X | Y | Total number of SNPs | Fraction | N’X’N’ | N’Y’N’ | X’ | Y’ | Total number of SNPs | Fraction | Difference (%) |
| | **Mouse SNPs** | | | | | | | | | | | | |
| 1 | CGA | CTA | G | T | 38698 | 0.26 | TAG | TCG | A | C | 38235 | 0.25 | 1.20 |
| 2 | AAG | ATG | A | T | 88927 | 0.59 | CAT | CTT | A | T | 89884 | 0.59 | 1.06 |
| 3 | CCC | CGC | C | G | 29680 | 0.20 | GCG | GGG | C | G | 29385 | 0.19 | 0.99 |
| 4 | CCT | CGT | C | G | 48777 | 0.32 | ACG | AGG | C | G | 49266 | 0.32 | 0.99 |
| 5 | AGA | ATA | G | T | 107141 | 0.71 | TAT | TCT | A | C | 108135 | 0.71 | 0.92 |
| 6 | GAC | GCC | A | C | 57560 | 0.38 | GGC | GTC | G | T | 57041 | 0.38 | 0.90 |
| 7 | TGA | TTA | G | T | 86691 | 0.57 | TAA | TCA | A | C | 87432 | 0.58 | 0.85 |
| 8 | GCT | GTT | C | T | 323720 | 2.14 | AAC | AGC | A | G | 326448 | 2.15 | 0.84 |
| 9 | AGC | ATC | G | T | 85493 | 0.56 | GAT | GCT | A | C | 84799 | 0.56 | 0.81 |
| 10 | CAA | CTA | A | T | 54492 | 0.36 | TAG | TTG | A | T | 54163 | 0.36 | 0.60 |
| 11 | TCG | TGG | C | G | 38824 | 0.26 | CCA | CGA | C | G | 39042 | 0.26 | 0.56 |
| 12 | GCT | GGT | C | G | 65256 | 0.43 | ACC | AGC | C | G | 64903 | 0.43 | 0.54 |
| 13 | TGG | TTG | G | T | 88219 | 0.58 | CAA | CCA | A | C | 88685 | 0.58 | 0.53 |
| 14 | GGA | GTA | G | T | 66022 | 0.44 | TAC | TCC | A | C | 66361 | 0.44 | 0.51 |
| 15 | TCT | TGT | C | G | 120710 | 0.80 | ACA | AGA | C | G | 121326 | 0.80 | 0.51 |
| 16 | TAC | TTC | A | T | 60192 | 0.40 | GAA | GTA | A | T | 60497 | 0.40 | 0.50 |
| 17 | AAA | ACA | A | C | 151052 | 1.00 | TGT | TTT | G | T | 150333 | 0.99 | 0.48 |
| 18 | CGT | CTT | G | T | 75731 | 0.50 | AAG | ACG | A | C | 75412 | 0.50 | 0.42 |

| | | | | | | | | | | | | |
|---|---|---|---|---|---|---|---|---|---|---|---|---|
| 19 | CCG | CTG | C | T | 348208 | 2.30 | CAG | CGG | A | G | 349621 | 2.31 | 0.40 |
| 20 | GAG | GCG | A | C | 50929 | 0.34 | CGC | CTC | G | T | 50724 | 0.33 | 0.40 |
| 21 | CAG | CCG | A | C | 56636 | 0.37 | CGG | CTG | G | T | 56410 | 0.37 | 0.40 |
| 22 | GAT | GTT | A | T | 63675 | 0.42 | AAC | ATC | A | T | 63432 | 0.42 | 0.38 |
| 23 | ACG | ATG | C | T | 480989 | 3.17 | CAT | CGT | A | G | 482775 | 3.18 | 0.37 |
| 24 | GGG | GTG | G | T | 74901 | 0.49 | CAC | CCC | A | C | 74624 | 0.49 | 0.37 |
| 25 | GCG | GTG | C | T | 276960 | 1.83 | CAC | CGC | A | G | 277835 | 1.83 | 0.31 |
| 26 | GAG | GGG | A | G | 225182 | 1.49 | CCC | CTC | C | T | 224605 | 1.48 | 0.26 |
| 27 | GAA | GCA | A | C | 95996 | 0.63 | TGC | TTC | G | T | 95759 | 0.63 | 0.25 |
| 28 | TCC | TGC | C | G | 56075 | 0.37 | GCA | GGA | C | G | 55960 | 0.37 | 0.21 |
| 29 | TCC | TTC | C | T | 351509 | 2.32 | GAA | GGA | A | G | 352103 | 2.32 | 0.17 |
| 30 | AAA | AGA | A | G | 362257 | 2.39 | TCT | TTT | C | T | 361808 | 2.39 | 0.12 |
| 31 | GAG | GTG | A | T | 73085 | 0.48 | CAC | CTC | A | T | 72996 | 0.48 | 0.12 |
| 32 | CCA | CTA | C | T | 283060 | 1.87 | TAG | TGG | A | G | 282755 | 1.87 | 0.11 |
| 33 | ACC | ATC | C | T | 262283 | 1.73 | GAT | GGT | A | G | 262546 | 1.73 | 0.10 |
| 34 | CAT | CCT | A | C | 83574 | 0.55 | AGG | ATG | G | T | 83647 | 0.55 | 0.09 |
| 35 | AGT | ATT | G | T | 100114 | 0.66 | AAT | ACT | A | C | 100033 | 0.66 | 0.08 |
| 36 | GGT | GTT | G | T | 95675 | 0.63 | AAC | ACC | A | C | 95744 | 0.63 | 0.07 |
| 37 | ACT | ATT | C | T | 384248 | 2.53 | AAT | AGT | A | G | 384516 | 2.54 | 0.07 |
| 38 | TAA | TGA | A | G | 285259 | 1.88 | TCA | TTA | C | T | 285425 | 1.88 | 0.06 |
| 39 | GAC | GGC | A | G | 253253 | 1.67 | GCC | GTC | C | T | 253112 | 1.67 | 0.06 |
| 40 | TAT | TGT | A | G | 477219 | 3.15 | ACA | ATA | C | T | 477431 | 3.15 | 0.04 |
| 41 | GCA | GTA | C | T | 255518 | 1.69 | TAC | TGC | A | G | 255481 | 1.69 | 0.01 |
| 42 | TAT | TTT | A | T | 103335 | 0.68 | AAA | ATA | A | T | 103323 | 0.68 | 0.01 |
| 43 | AAG | AGG | A | G | 299963 | 1.98 | CCT | CTT | C | T | 299979 | 1.98 | 0.01 |
| 44 | CAA | CGA | A | G | 264331 | 1.74 | TCG | TTG | C | T | 264323 | 1.74 | 0.00 |
| 45 | TAA | TTA | A | T | 133002 | 0.88 | | | | | | | |
| 46 | CAG | CTG | A | T | 84745 | 0.56 | | | | | | | |
| 47 | GCC | GGC | C | G | 49385 | 0.33 | | | | | | | |
| 48 | ACT | AGT | C | G | 107564 | 0.71 | | | | | | | |
| 49 | CCG | CGG | C | G | 34929 | 0.23 | | | | | | | |
| 50 | AAT | ATT | A | T | 122244 | 0.81 | | | | | | | |
| 51 | TCA | TGA | C | G | 65934 | 0.43 | | | | | | | |
| 52 | GAC | GTC | A | T | 52841 | 0.35 | | | | | | | |

**Supplementary Table S2**. Mutation spectra inferred from single nucleotide insertion or deletion (indels) in the genomes in human populations.

| N° | N Δ N | N X N | Δ | X | Total number of indels | Fraction | N' Δ N' | N'X'N' | Δ | X' | Total number of indels | Fraction | Difference (%) |
|---|---|---|---|---|---|---|---|---|---|---|---|---|---|
| | Triplets with sequence context in which a base corresponding to indel is different from the neighboring base. | | | | | | | | | | | | |
| 1 | C-G | CTG | - | T | 31093 | 1.74 | C-G | CAG | - | A | 31132 | 1.75 | 0.13 |
| 2 | G-C | GTC | - | T | 6049 | 0.34 | G-C | GAC | - | A | 6092 | 0.34 | 0.71 |
| 3 | T-C | TGC | - | G | 11496 | 0.64 | G-A | GCA | - | C | 11346 | 0.64 | 1.30 |
| 4 | T-T | TAT | - | A | 33846 | 1.90 | A-A | ATA | - | T | 33349 | 1.87 | 1.47 |
| 5 | T-T | TGT | - | G | 44135 | 2.48 | A-A | ACA | - | C | 43138 | 2.42 | 2.26 |
| 6 | T-C | TAC | - | A | 7640 | 0.43 | G-A | GTA | - | T | 7449 | 0.42 | 2.50 |
| 7 | C-A | CTA | - | T | 16203 | 0.91 | T-G | TAG | - | A | 15787 | 0.89 | 2.57 |
| 8 | A-C | ATC | - | T | 11841 | 0.66 | G-T | GAT | - | A | 11511 | 0.65 | 2.79 |
| 9 | A-T | AGT | - | G | 22001 | 1.23 | A-T | ACT | - | C | 22719 | 1.27 | 3.16 |
| 10 | A-G | ATG | - | T | 17041 | 0.96 | C-T | CAT | - | A | 16374 | 0.92 | 3.91 |
| 11 | G-T | GCT | - | C | 15314 | 0.86 | A-C | AGC | - | G | 14649 | 0.82 | 4.34 |
| 12 | A-A | AGA | - | G | 36802 | 2.06 | T-T | TCT | - | C | 38617 | 2.17 | 4.70 |
| 13 | G-G | GAG | - | A | 16611 | 0.93 | C-C | CTC | - | T | 17504 | 0.98 | 5.10 |
| 14 | C-C | CGC | - | G | 2900 | 0.16 | G-G | GCG | - | C | 3063 | 0.17 | 5.32 |
| 15 | C-C | CAC | - | A | 13485 | 0.76 | G-G | GTG | - | T | 12680 | 0.71 | 5.97 |
| 16 | A-G | ACG | - | C | 4610 | 0.26 | C-T | CGT | - | G | 4249 | 0.24 | 7.83 |
| 17 | T-G | TCG | - | C | 2880 | 0.16 | C-A | CGA | - | G | 2603 | 0.15 | 9.62 |
| 18 | T-A | TGA | - | G | 16897 | 0.95 | T-A | TCA | - | C | 19855 | 1.11 | 14.90 |
| | Triplets with sequence context in which a base corresponding to indel is the same as neighboring base. | | | | | | | | | | | | |
| 19 | A-A | AAA | - | A | 19326 | 1.08 | T-T | TTT | - | T | 19540 | 1.10 | 1.10 |
| 20 | G-G | GGG | - | G | 1144 | 0.06 | C-C | CCC | - | C | 1124 | 0.06 | 1.75 |
| 21 | T-G | TTG | - | T | 11150 | 0.63 | C-A | CAA | - | A | 123998 | 6.95 | 91.01 |
| 22 | C-T | CTT | - | T | 138611 | 7.77 | A-G | AAG | - | A | 11140 | 0.62 | 91.96 |
| 23 | A-C | AAC | - | A | 4327 | 0.24 | G-T | GTT | - | T | 79624 | 4.47 | 94.57 |
| 24 | T-A | TTA | - | T | 6518 | 0.37 | T-A | TAA | - | A | 133014 | 7.46 | 95.10 |
| 25 | A-T | ATT | - | T | 155454 | 8.72 | A-T | AAT | - | A | 7175 | 0.40 | 95.38 |
| 26 | T-C | TTC | - | T | 5072 | 0.28 | G-A | GAA | - | A | 112956 | 6.34 | 95.51 |
| 27 | C-G | CCG | - | C | 337 | 0.02 | C-G | CGG | - | G | 14700 | 0.82 | 97.71 |
| 28 | G-T | GGT | - | G | 1290 | 0.07 | A-C | ACC | - | C | 58855 | 3.30 | 97.81 |
| 29 | T-C | TCC | - | C | 62091 | 3.48 | G-A | GGA | - | G | 1287 | 0.07 | 97.93 |
| 30 | T-G | TGG | - | G | 73594 | 4.13 | C-A | CCA | - | C | 1510 | 0.08 | 97.95 |
| 31 | C-T | CCT | - | C | 1435 | 0.08 | A-G | AGG | - | G | 72608 | 4.07 | 98.02 |
| 32 | G-C | GCC | - | C | 41325 | 2.32 | G-C | GGC | - | G | 750 | 0.04 | 98.19 |
